# Random recurrent networks near criticality capture the broadband power distribution of human ECoG dynamics

**DOI:** 10.1101/036228

**Authors:** Rishidev Chaudhuri, Biyu J. He, Xiao-Jing Wang

## Abstract

The power spectrum of brain electric field potential recordings is dominated by an arrhythmic broadband signal but a mechanistic account of its underlying neural network dynamics is lacking. Here we show how the broadband power spectrum of field potential recordings can be explained by a simple random network of nodes near criticality. Such a recurrent network produces activity with a combination of a fast and a slow autocorrelation time constant, with the fast mode corresponding to local dynamics and the slow mode resulting from recurrent excitatory connections across the network. These modes are combined to produce a power spectrum similar to that observed in human intracranial EEG (i.e., electrocorticography, ECoG) recordings. Moreover, such a network naturally converts input correlations across nodes into temporal autocorrelation of the network activity. Consequently, increased independence between nodes results in a reduction in low-frequency power, which offers a possible explanation for observed changes in ECoG power spectra during task performance. Lastly, changes in network coupling produce changes in network activity power spectra reminiscent of those seen in human ECoG recordings across different arousal states. This model thus links macroscopic features of the empirical ECoG power spectrum to a parsimonious underlying network structure and proposes potential mechanisms for changes in ECoG power spectra observed across behavioral and arousal states. This provides a computational framework within which to generate and test hypotheses about the cellular and network mechanisms underlying whole brain electrical dynamics, their variations across behavioral states as well as abnormalities associated with brain diseases.

## Acknowledgments

This work was supported by the Office of Naval Research Grant N00014-13-1-0297, National Institutes of Health Grant R01 MH062349 (to X.J.W.) and by the intramural research program of the National Institutes of Health / National Institute of Neurological Disorders and Stroke (to B.J.H.). We thank Alberto Bernacchia, John Murray and Guangyu Yang for discussions. The authors declare no competing financial interests.

## Introduction

The power spectrum of electrical field potentials recorded from the brain consists of a set of oscillatory peaks, indicative of underlying rhythmicity, riding on top of a broadband “1 */f*^*β*^” slope (power falls off with frequency, following *P ≈ A/f^β^*, where *β* is the power-law exponent), which constitutes the majority of signal power. Research over the past decades has significantly advanced our understanding of the functional roles and generative mechanisms of brain oscillations at different frequencies (Buzsaki, 2006; Fries, 2009; Wang, 2010; Jensen et al., 2012; Womelsdorf et al., 2014). However, the origins of the arrhythmic signal contributing the 1/*f^β^* component of the spectrum remain elusive (Bedard and Destexhe, 2009; Roberts et al., 2015).

Recent research has shown that this arrhythmic, broadband field potential cannot be explained as summation of many oscillations (Miller et al., 2009b; He et al., 2010). By contrast, it appears to be a distinct type of brain activity, potentially a macroscopic manifestation of the irregular firing of cortical neurons (Miller et al., 2009a; He, 2014). In particular, broadband power in the gamma-frequency (>30 Hz) range tightly correlates with population firing rate (Manning et al., 2009; Whittingstall and Logothetis, 2009; Ray and Maunsell, 2011; Buzsáki and Wang, 2012) and exhibits functional specificity across a variety of tasks (Crone et al., 1998; Miller et al., 2009b; Ossandón et al., 2011; Bouchard et al., 2013). In the very low frequency range (<1 Hz), the slope of the power spectrum (i.e., the power-law exponent *β*) – an index of the amount of long-time autocorrelation in the signal – is reduced during a visual detection task (He et al., 2010). Despite these results demonstrating the functional significance of arrhythmic, broadband activity, a mechanistic account explaining the full frequency range of its signature power spectrum remains lacking.

Multiple studies using local field potential (LFP) and ECoG recordings have found the power law exponent *β* is typically between 2 and 3 (Milstein et al., 2009; Miller et al., 2009a; Manning et al., 2009; Freeman and Zhai, 2009; He et al., 2010). A study using DC-recording revealed the shape of human ECoG power spectra across a wide range of frequencies, from 0.01 Hz to >100 Hz (He et al., 2010). The power spectra exhibit a distinctive shape: at very low frequencies (<0.1 Hz) and above 1 Hz, power scales approximately proportional to the inverse-square of frequency (*P* ∼ 1/*f*^2^), while power spectra in the intermediate frequency range (0.1-1 Hz) are much flatter. This tripartite shape was conserved across subjects, albeit with differences in the locations of the transitions between the decaying and flat regions of the spectrum.

Here, we use a network model to investigate the potential neural-circuit-level origins of the broadband signal in field potential recordings. We find that the power spectrum is well fit as the combination of two linear modes, which sum to produce the characteristic tripartite shape of the empirically-observed human ECoG spectrum. We then show that such a power spectrum generically emerges from the activity of a recurrent network model with nodes randomly connected to each other, provided that the net excitation (i.e. excess of excitation over inhibition) between nodes roughly balances the intrinsic decay of activity. In this sense, the network is close to dynamical criticality (Beggs and Timme, 2012; Deco and Jirsa, 2012; Priesemann et al., 2014; Roberts et al., 2015; Bellay et al., 2015). We characterize the dependence of the power spectrum on network parameters and on input structure, and show that such random recurrent networks naturally convert spatial correlation in the input into temporal correlations in network activity. We then extend the architecture and investigate networks with a distance-dependent connectivity profile and networks where the nodes are themselves clusters of sub-nodes. Our analyses link empirically observed human ECoG power spectra to plausible underlying neural network dynamics and suggest potential circuit-level explanations for changes in the low-frequency power spectrum across behavioral and arousal states.

## Results

### Two modes in the low-frequency power spectrum

The power-law exponents seen in ECoG and LFP power spectra (Fig. 1A) are characteristic of linear systems, which have an autocorrelation function composed of a weighted sum of exponentials. The power spectrum of an exponential function, *e^−λt^*, is proportional to 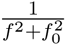, where *f* is the frequency and *f*_0_ = *λ*/2*π*. As shown in Fig. 1B, when plotted on a log-log scale these functions (often called Lorentzians) are approximately flat at low frequencies and scale with slope –2 at high frequencies; the transition between the two regimes happens at the frequency *f*_0_, which we refer to as the “knee” frequency.

**Figure 1.**
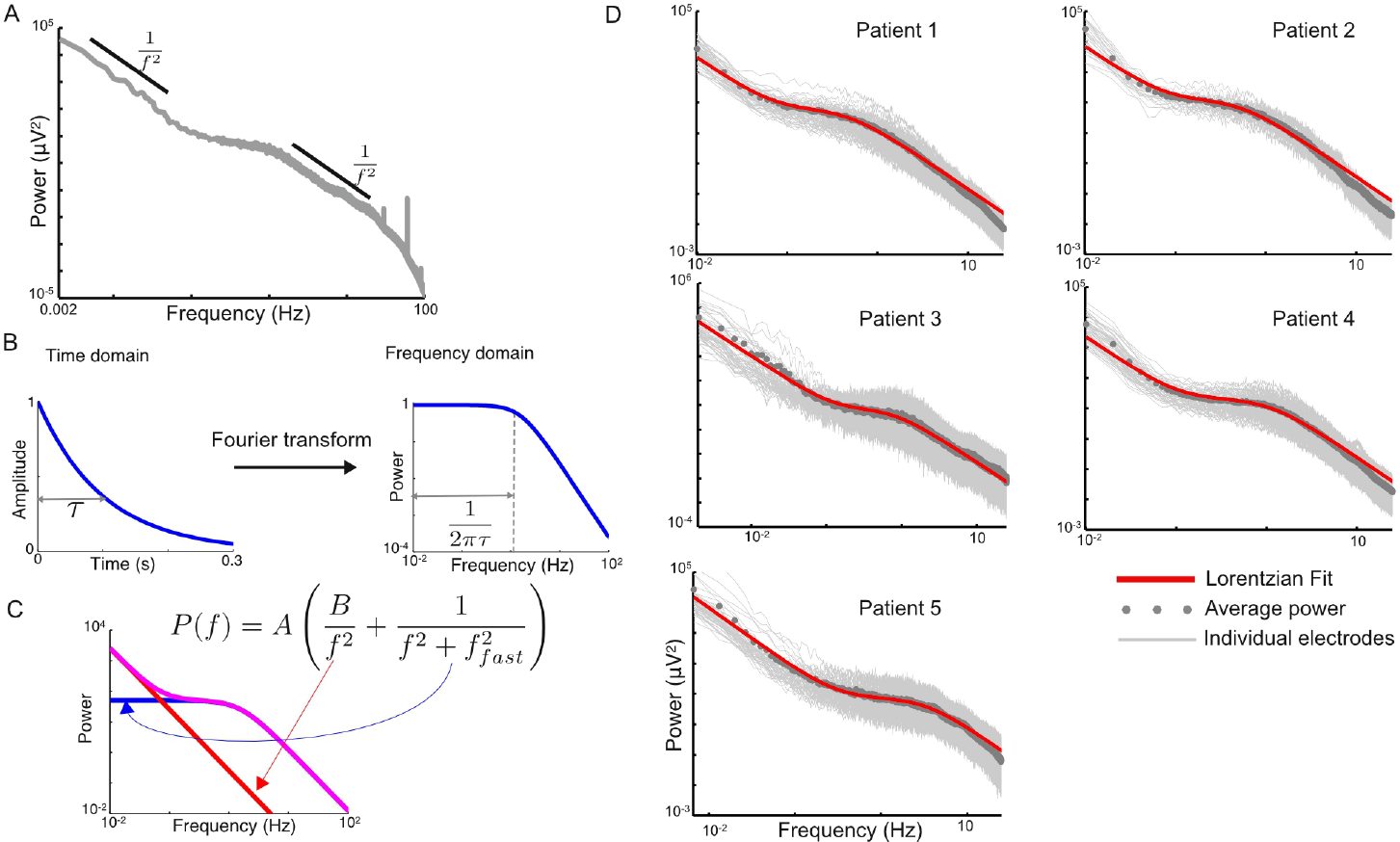
The low-frequency power spectra of human ECoG are well-fit by the sum of two Lorentzian functions. **A**, Average power spectrum from one patient in the study of He et al. (2010). The power spectrum resembles a power law with frequency dependence 1/*f*^2^ at low and high frequencies, with a roughly flat intermediate region. **B**, The power spectrum of an exponential is a Lorentzian function, which shows near-flat behavior at low-frequencies and 1/*f*^2^ scaling at high frequencies, with a transition point set by the time-constant of the exponential. **C**, The sum of two Lorentzian functions yields a shape resembling the power spectrum of Fig. 1A, with the “knee” frequency set by *f_fast_*. **D**, Each plot is the power spectrum from one patient in He et al. (2010). The light grey traces correspond to recordings from each electrode while the dark grey circles are the averages across all electrodes. Red traces are fits of a sum of two Lorentzian functions (corresponding to the functional form shown in Fig. 1C). The functional form is fit to the frequencies below 5 Hz and the data is shown up to 25 Hz. The slope of the power spectrum is steeper for frequencies beyond 25 Hz; see Discussion and Fig. 8 for fits to the remainder of the spectrum.

Motivated by this observation, we fit the power spectrum from the 5 subjects of He et al. (2010) as the weighted sum of a fast and a slow linear mode (Fig. 1C). The corresponding functional form is

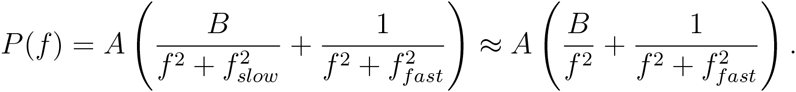

In the second equation we have assumed that *f_slow_* is small enough to be outside the observational range and thus can be neglected. Thus, the fit with two Lorentzians has a single knee frequency, located at *f_fast_*.

In Fig. 1D we show this fit for the average power spectrum of each of the 5 subjects in the study of He et al. (2010). The functional form accounts for the shape of the power spectrum across several orders of magnitude (with deviations at high frequencies; see Fig. 8). The location of the transition from the initial 1/*f*^2^ behavior to the flat region is set by the relative contributions of the two Lorentzians and hence is determined by the parameter B. As previously mentioned, the intermediate transition to 1/*f*^2^ has location set by *f_fast_*. For the five patients shown in Fig. 1C, the knee frequency (*f_fast_*) is at 0.49 Hz, 0.55 Hz, 0.81 Hz, 1.10 Hz and 3.47 Hz respectively.

In Fig. 2, we show the knee frequency (*f_fast_*) for individual fits to each electrode in each patient. As can be seen, there is considerable variation in the characteristic frequency across electrodes, with the fastest frequency being 2-5 times the slowest one. However, neighboring electrodes show similar values for the knee frequency, with correlations of 0.35, 0.31, 0.28, 0.32, and 0.33 respectively (p<0.002 for all patients; see Methods for further details). This suggests that the variation is not random and contains spatial structure (see Discussion).

**Figure 2.**
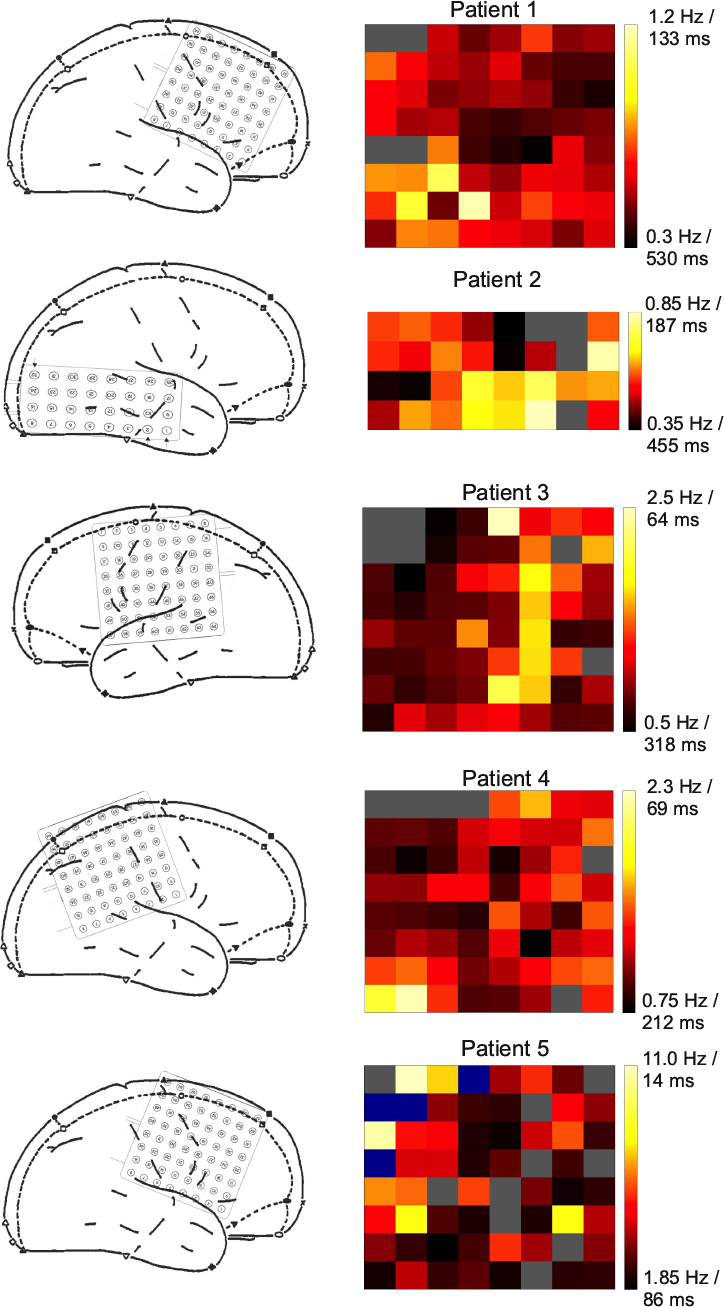
The knee frequency (*f_fast_)* for individual electrodes across patients. Left panel: Locations of electrodes for each of the five patients. Right panel: knee frequency for individual electrodes, with location in the heat map corresponding to the electrode locations shown on the left. Electrodes excluded in the study of He et al. (2010) are shown in grey and, for Patient 5, electrodes poorly fit as a sum of Lorentzians are shown in blue. Timescales shown are the time-constants of a linear system with corresponding knee frequency (i.e. *τ* = 1/2π*_knee_*).

### A random network model for the power spectrum

We next construct a recurrent network model which reproduces the observed power spectrum. The model network has *N* nodes, which could be neurons or networks of neurons. The *j*th node has activity *r_j_*, which evolves in time according to the equation

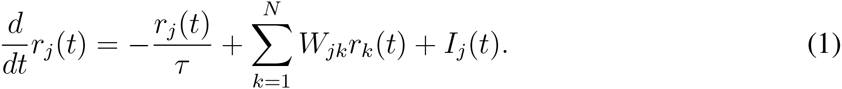

Each node receives input from the other nodes in the network (*r_k_*) with weight *W_jk_* and also receives some external input *I_j_*, corresponding to input that does not come from within the network. In the absence of any input, the firing rate of the j th node decays exponentially to 0 with a rate given by *τ*. Grouping the firing rates into a vector and the weights into a matrix, we can rewrite this equation as

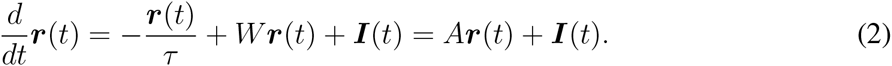

Here the matrix *A* is identical to *W* except along the diagonal, where we subtract 1/*τ* to account for the intrinsic decay of activity.

We assume that the observed field potential recording results from summing together the activity of a subset of network nodes. Thus, if *x(t)* is this summed activity, we have

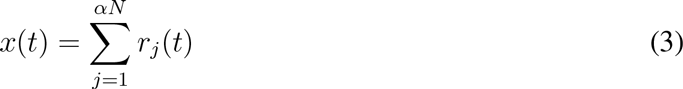

Here 0 < *α* < 1 is the fraction of the network we are summing over, and we have written the sum over the first *αN* nodes for convenience. We will refer to *x(t)* as “network activity”. While we consider an equally-weighted sum of nodes contributing to *x*, our analysis can easily be extended to a differentially-weighted sum or spatial kernel applied to the nodes.

We choose the connections (the entries of matrix *W*) to be sparse and random: each entry is non-zero with probability *p*, and non-zero entries are drawn from a normal distribution: *W_ij_* ∼ 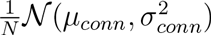. In Fig. 3B, we show the power spectrum of network activity. This reproduces the data for appropriately chosen values of *τ, μ_conn_* and *α*. Note that *μ_conn_* must be set to almost balance the intrinsic decay of network activity (which decays with time-constant τ at each node).

**Figure 3.**
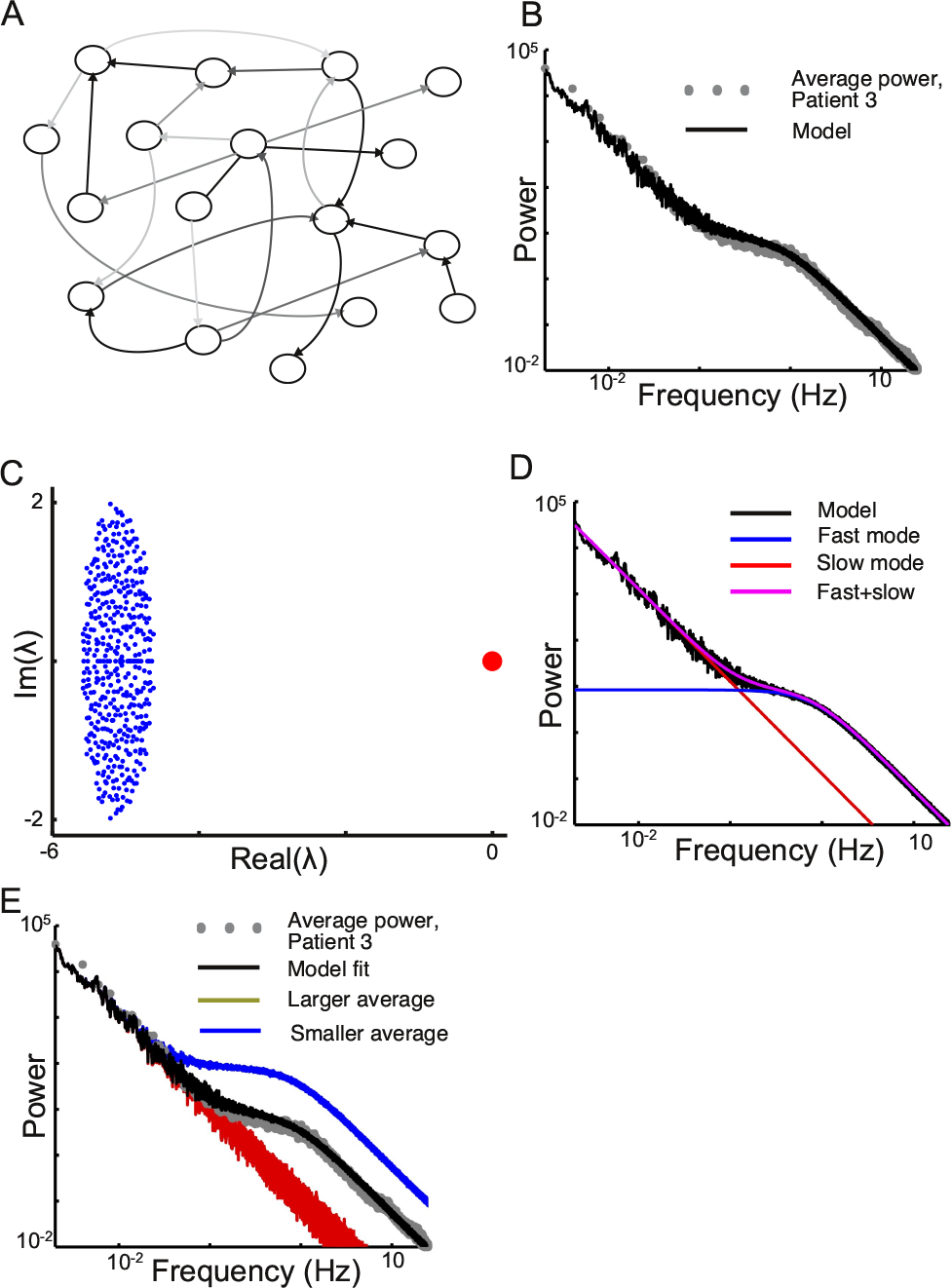
The power spectrum of a random network reproduces observed ECoG power spectra. **A**, Schematic of a sparse randomly-connected network. **B**, Power spectrum of network activity in a random network where mean connection strengths approximately balance the intrinsic decay of activity. The power spectrum from patient 3 is shown for comparison. **C**, The eigenvalue spectrum of the network coupling matrix shows a cluster of fast modes (in blue) and a single slow mode (in red). **D**, The power spectrum of the simulated network is the sum (purple) of Lorentzians contributed by the fast modes (blue) and the slow mode (red). **E**, Effect of spatial averaging on power spectrum. Black: power spectrum from the network in panel B with the same degree of averaging. Blue: network activity derived by summing over a smaller fraction of the network (here, a single node). Red: network activity derived by summing over a larger fraction of the network (here, all nodes in the network). The slow mode is shared between nodes, while the fast modes are uncorrelated; thus averaging over nodes boosts the contribution of the slow mode. In particular, summing over the entire network yields a power spectrum that shows pure power-law behavior.

The activity of the multidimensional linear system in Eq. 2 can be thought of geometrically: the vector *r* lives in an *N*-dimensional space with each dimension corresponding to the activity at one node (i.e., *r_j_* is the activity along the *j*th node). The system can be solved by changing the coordinate system and rewriting the activity vector, *r*, in a new coordinate system whose directions are given by the eigenvectors of the matrix *A*. These eigenvectors form the natural coordinate system in which to see the activity of *A*: they provide a decomposition of the network activity into a set of characteristic modes, each of which evolves independently in time with its own characteristic timescale. This decouples an *N*-dimensional problem into a collection of *N* one-dimensional problems.

The eigenvectors are defined as vectors *v_m_* that satisfy the equation *Av_m_ =λ_m_v_m_*, where *λ_m_* is a constant, called the eigenvalue corresponding to the eigenvector *v_m_*. The characteristic timescale of network activity corresponding to the eigenvector *v_m_* is – 1/ ℜe(λ_m_). Thus, the eigenvalues tell us what timescales the network will show, and the eigenvectors tell us how this activity is distributed across nodes.

To understand how the network model is able to reproduce the data, we consider the distribution of eigenvalues of the network coupling matrix, *A*. These eigenvalues describe the characteristic temporal modes of the network (see Methods for more details). For the randomly-connected network we consider, the eigenvalues take a particularly simple form, known from the theory of random matrices (Rajan and Abbott, 2006; Ganguli et al., 2008; Tao, 2011) and depicted in Fig. 3C. The network has one slow mode, here corresponding to an eigenvalue near 0 (the red point in Fig. 3C), and *N* – 1 fast modes, which are centered around – 1/τ (the cloud of blue points in Fig. 3C). If the external input to each node is independent, then the power spectrum of network activity can be approximately broken up into contributions from each of these two sets of modes (Fig. 3D and see Methods for derivation), and is given by

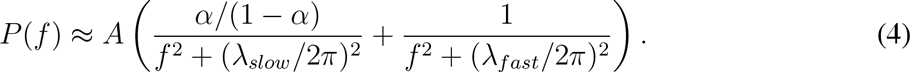

Thus, if *λ_slow_* is small (i.e. the corresponding mode is very slow), the power spectrum is of the same form as the fit in Fig. 1, with the relative contribution of the slow and fast modes determined by *α* (the fraction of the network averaged over), and the location of the knee frequency given by *λ_fast_*/2*π*.

The eigenvalue *λ_slow_* emerges from inter-node recurrent interactions: in response to an input, nodes of the network excite each other, thus reverberating the signal around the network and slowing its decay. *λ_slow_* is approximately located at *μ – 1/τ*, where *μ = μ_conn_ρ*, and when the recurrent excitation (*μ*) balances the intrinsic decay of activity (1/*τ*) this is very close to 0 (see Methods). In this case signals reverberate in the network for a long time, giving rise to a slowly-decaying autocorrelation function.

On the other hand, *λ_fast_* is set by the intrinsic properties of each network node and is approximately located at – 1/*τ*. There are *N* – 1 such fast modes, each with time-constant approximately equal to *τ*. Note that if each node is a cluster of neurons, the time-constant *τ* itself emerges from underlying recurrent interactions; we return to this issue later.

Since the slow mode emerges from inter-node interactions, it corresponds to a spatially-distributed pattern of network activity. This is given by the eigenvector, *v_slow_*, corresponding to the eigenvalue *λ_slow_*. *v_slow_* can be thought of as a slowly-varying background state of the network that all the nodes are coupled to; in particular, for low variability in connection strengths, *v_slow_* has approximately equal weight at each node. By contrast, the fast modes are uncorrelated with each other and thus different nodes participate in a particular fast mode to greatly varying degrees. As a consequence of this spatial distribution of the fast and slow modes, the network model accounts for the common observation that low frequency activity (i.e. slow timescales) shows a wider spatial correlation than activity at high frequencies (Leopold et al., 2003; von Stein and Sarnthein, 2000; Buzsaki, 2006).

Moreover, because of the broader spatial distribution of the slow mode, averaging activity across multiple nodes in a network will emphasize the slow mode and increase its contribution to the observed power spectrum. This can be seen in the equation for the power spectrum of network activity above, where the relative contributions of the slow and fast modes are given by *α*, the fraction of the network averaged over to generate the network activity. In Fig. 3E, we show the effect of averaging over different fractions of the network. In particular, averaging over a large fraction of nodes yields a power spectrum dominated by the 1/*f*^2^ term.

Note, however, that if there exist multiple recurrent networks, each described by an equation such as Eq. 1 but only weakly coupled to each other, then averaging across nodes belonging to these different networks will not change the shape of the power spectrum, because the weak coupling between these networks would not generate another slow mode.

While the network model has purely excitatory connections between the nodes (recall that these nodes may, in turn, be clusters of neurons), a similar picture holds for a network where some fraction of the nodes have inhibitory projections onto their targets and others have excitatory projections (Fig. 4A). As previously argued (Rajan and Abbott, 2006; Ganguli et al., 2008), and as depicted in Fig. 4B, a randomly-connected network where a subset of nodes make inhibitory connections onto their targets has an eigenvalue spectrum that is similar to that of Fig. 3. Here the location of the slow network mode depends on the difference between excitation and inhibition. The slow eigenvalue is located at *μ_E_ρ_E_* – *μ_I_p_I_* – 1/*τ*, where *μ_Ε_* and *μ_I_* are the magnitudes of the coupling strengths for excitation and inhibition respectively, *p_E_* and *p_I_* are the respective connection probabilities, and, as before, *τ* is the intrinsic decay time-constant (see Methods). If this difference between excitation and inhibition closely balances the intrinsic decay then the network will show long timescales. In Fig. 4C we show how the power spectrum of a network with 80% excitatory and 20% inhibitory connections can reproduce the observed power spectra.

**Figure 4.**
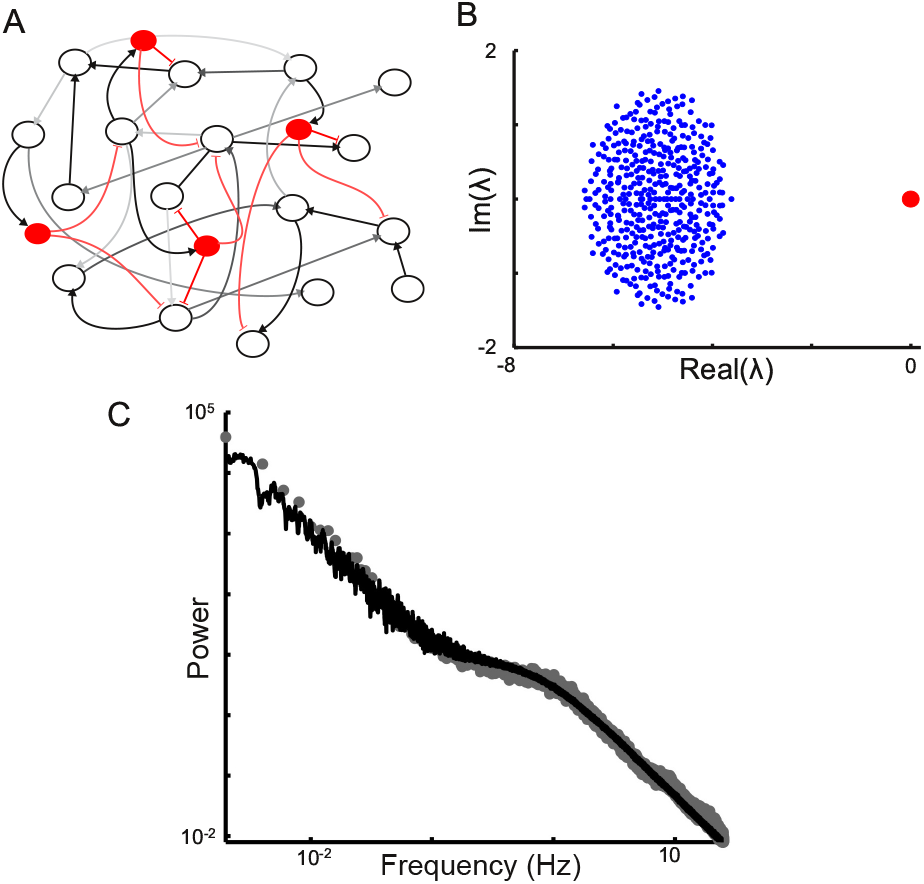
Power spectrum of a random network with both excitatory and inhibitory connections. **A**, Schematic of network, as in Fig. 3A but with the incorporation of interneurons shown in red. **B**, Eigenvalue spectrum of the network. **C**, Power spectrum of network activity (black) along with data from patient 3 (grey filled circles).

The low frequency component of the ECoG power spectrum has been shown to differ between arousal states (i.e. waking vs sleep) and to change upon task initiation (He et al., 2010). We next investigate manipulations of the model that may underlie such state-dependent changes in the shape of the low-frequency power spectrum.

## Correlations in the input preferentially drive slow timescales

The eigenvector corresponding to a particular timescale determines not only the spatial distribution of that mode, but also how much that mode is activated in response to a given profile of input. Given a particular spatial pattern of input, the correlation of this spatial pattern with an eigenvector determines how strongly the corresponding temporal mode is driven (this is approximately true, but see Eq. M6 in Methods for a more precise statement). This corresponds to the intuition that input whose spatial distribution resembles a particular eigenvector should preferentially activate the temporal mode corresponding to that eigenvector.

The slow and fast modes have different spatial distributions, and thus are differentially driven by various inputs (see Eq. M14 for the power spectrum). Since the slow mode is shared across the network, it is driven by the component of the input that is common between nodes. By contrast, input that is uncorrelated between nodes drives the slow network mode incoherently, with some nodes contributing positively and others negatively, so that the net effect is small. As a consequence, a random recurrent network architecture generically transforms correlations in input across nodes into long temporal correlations in network activity. In Fig. 5 we show how the power spectrum of node activity depends on the degree of spatial correlation in the input (recall that the power spectra in Fig. 3 are for uncorrelated input). In particular, we note that a spatial decorrelation of input to a network would lead to a reduction in low-frequency power, as observed in ECoG power upon task-initiation (He et al., 2010).

**Figure 5.**
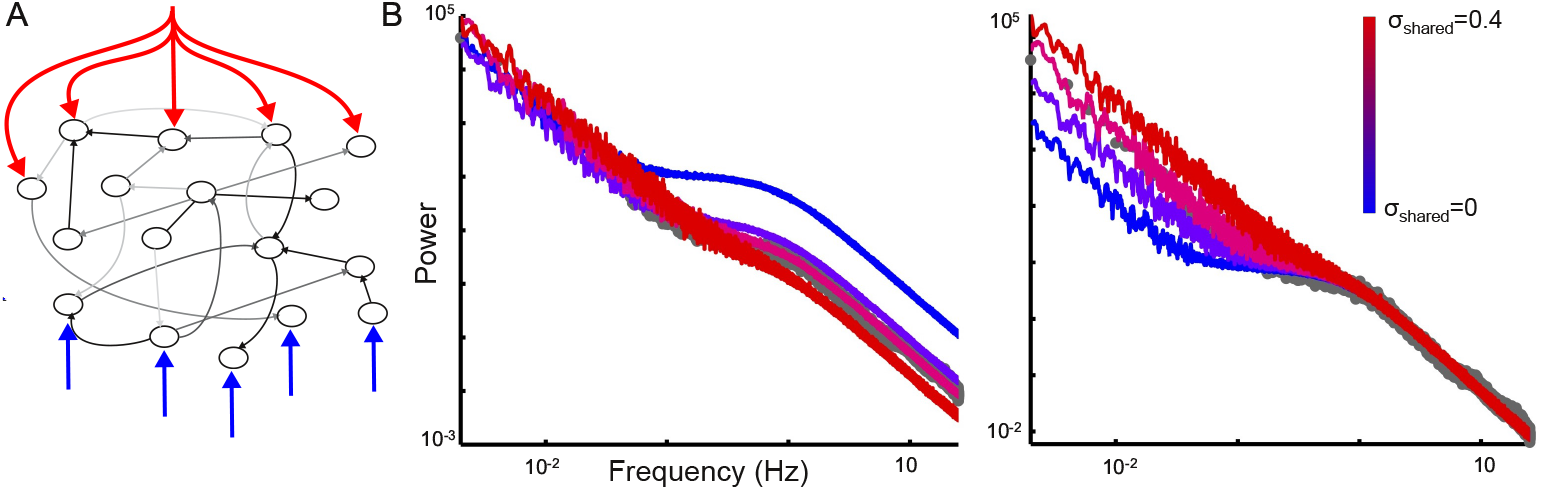
The network converts shared input into network activity with long temporal correlations. (A) Schematic of a random network with nodes that receive both shared input (red) and uncorrelated input (blue). (B) Power spectra for network activity in response to different fractions of correlated input. Left panel shows normalized power spectra, while the right panel highlights that correlated input leads to an increase in low-frequency power in the network. The average power spectrum from Patient 3 is shown as dark gray circles, for comparison. To highlight the role of correlated input in driving the slow mode, we average over a smaller fraction of network nodes. Thus, the blue trace (without input correlations) reflects the fast modes to a greater extent than the data, but this can be compensated for by more correlation in the input. *σ_shared_* is the variance of the common input; the variance of the remaining input (uncorrelated across nodes) is chosen so that total variance is constant (see Methods). Note that the power spectrum is still well-fit by the sum of two Lorentzians, with the amplitudes depending on the degree of correlation (Eq. M14).

### Distance-dependent connection probability changes the slope of low-frequency power spectrum

We have assumed that the nodes in the network are connected to each other with equal probability and mean weight. However, networks of neurons that are widely-distributed in space typically have a distance-dependence in connection probability and number: several studies have concluded that neural connectivity is primarily local and, despite notable exceptions, tends to decay with distance both within a cortical area and between cortical areas (Destexhe et al., 1999; Ercsey-Ravasz et al., 2013; Markov et al., 2014). We now consider model networks with nodes that have some underlying spatial location and whose connection strength decays exponentially with distance.

In Fig. 6A, we show the eigenvalue distributions of three such networks with progressively sharper connectivity profiles. In contrast to the completely random network of Fig. 3, these networks contain a number of intermediate eigenvalues between the slow shared mode and the cluster of modes around the single-node timescale. As the decay of connections with distance becomes sharper, the number of intermediate eigenvalues increases; this can be understood by noting that the positions of the eigenvalues are approximately given by the Fourier transform of the connectivity profile (see Methods), and hence networks with sharply-localized connectivity will have eigenvalues that are more spread out.

**Figure 6.**
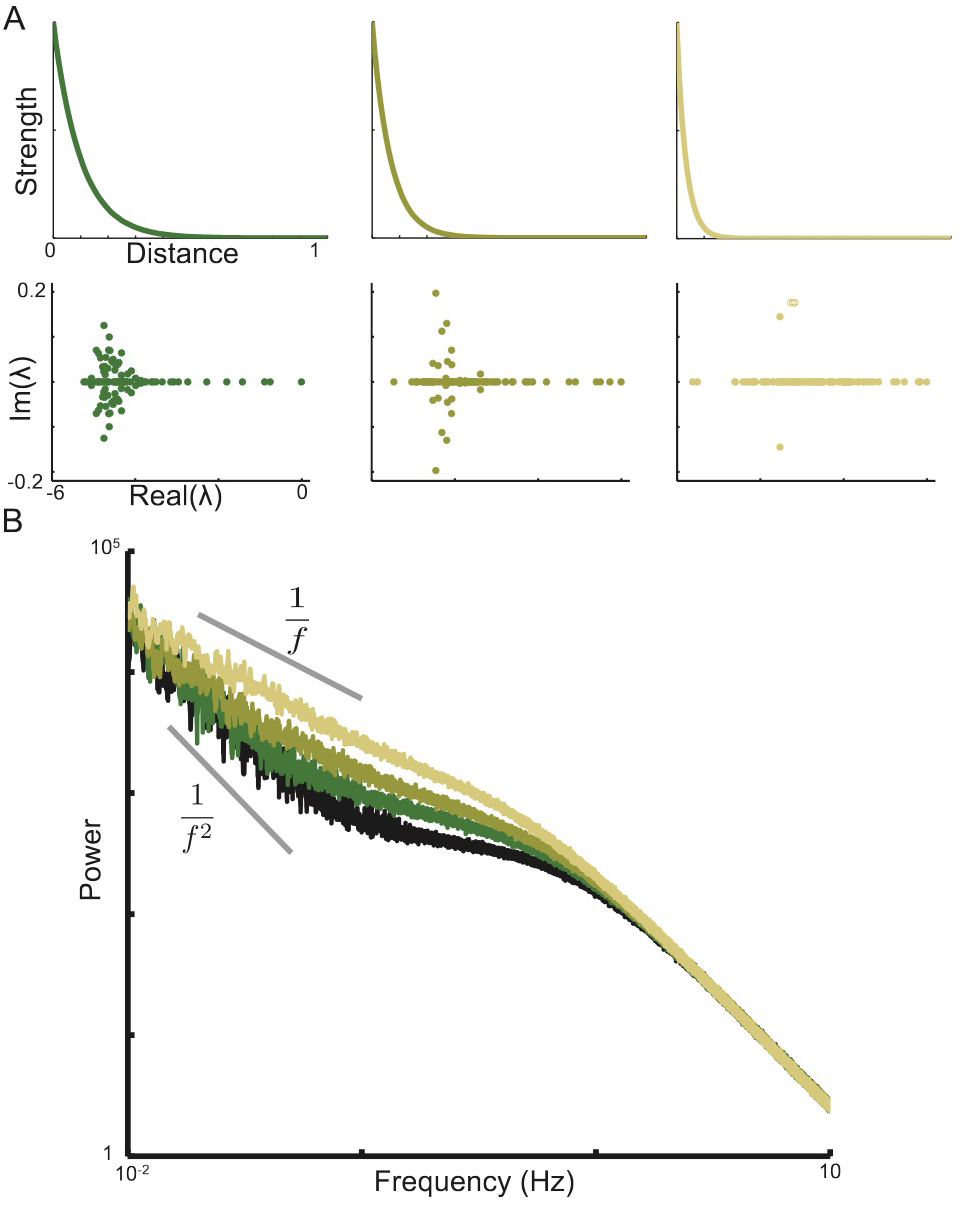
Network endowed with distance-dependent connectivity yields shallower power spectra. **A**, Connectivity profile of three networks with increasingly sharp decay of connection strength with distance (top), and the corresponding eigenvalue spectra (bottom). **B**, Power spectra for the three networks shown in A (in successively lighter shades of green) along with the power spectrum from Fig. 3, for comparison (shown in black).

In Fig. 6B we show the effect of distance-dependent connectivity on the network power spectram. Heuristically, the intermediate eigenvalues contribute Lorentzian functions with knee frequencies located in the low-frequency part of the power spectrum. These combine to produce a shallower low-frequency slope. As the distance-dependent decay of connectivity becomes steeper there are more such intervening modes, and the slope of the low-frequency network power spectrum continues to become shallower, leading to a scaling of power with frequency that is closer to 1/*f* (as seen in the light green trace of Fig. 6B). Thus networks with predominantly local connectivity could underlie experimental observations of 1/*f* power spectra in recordings (Bedard et al., 2006; Bedard and Destexhe, 2009; Dehghani et al., 2010), and different distance-dependent profiles of connectivity could explain differences in the slope of low-frequency power spectra between subjects, regions of the cortex or arousal states.

### Clustered network architectures

Thus far we have treated the nodes in our network as single entities with no internal structure. While it is possible that the nodes correspond to single neurons, the node timescales we observe are on the order of hundreds of milliseconds. This is longer than membrane time constants and most synaptic time constants; however it is within the range of other long cellular time constants (Carter and Wang, 2007; Zhang and Seguela, 2010; Letzkus et al., 2011), and we consider these further in the Discussion. An alternative hypothesis is that the nodes correspond to cell assemblies or clusters of neurons, with the individual neurons in these clusters showing faster timescales (on the order of milliseconds) and the timescales of each cluster emerging from inter-neuron recurrent interactions. The resulting architecture would thus be hierarchical: individual neurons form clusters via recurrent interactions and the clusters further interact to produce the very long network timescales.

The eigenvalue spectrum of a network with such a clustered structure is shown in Fig. 7B. For a network with N clusters, the eigenvalue spectrum shows a single slow mode near 0 (red circle) and *N* – 1 faster modes distributed around the time-constant of a single cluster (blue circles). Thus, the long timescale behavior is the same as before. However, the eigenvalue spectrum also shows a number of much faster modes clustered around the intrinsic time-constant of a single neuron (black circles). In the lower panel of Fig. 7B we highlight these two regions of the eigenvalue spectrum, revealing the signature of the underlying hierarchical architecture.

**Figure 7.**
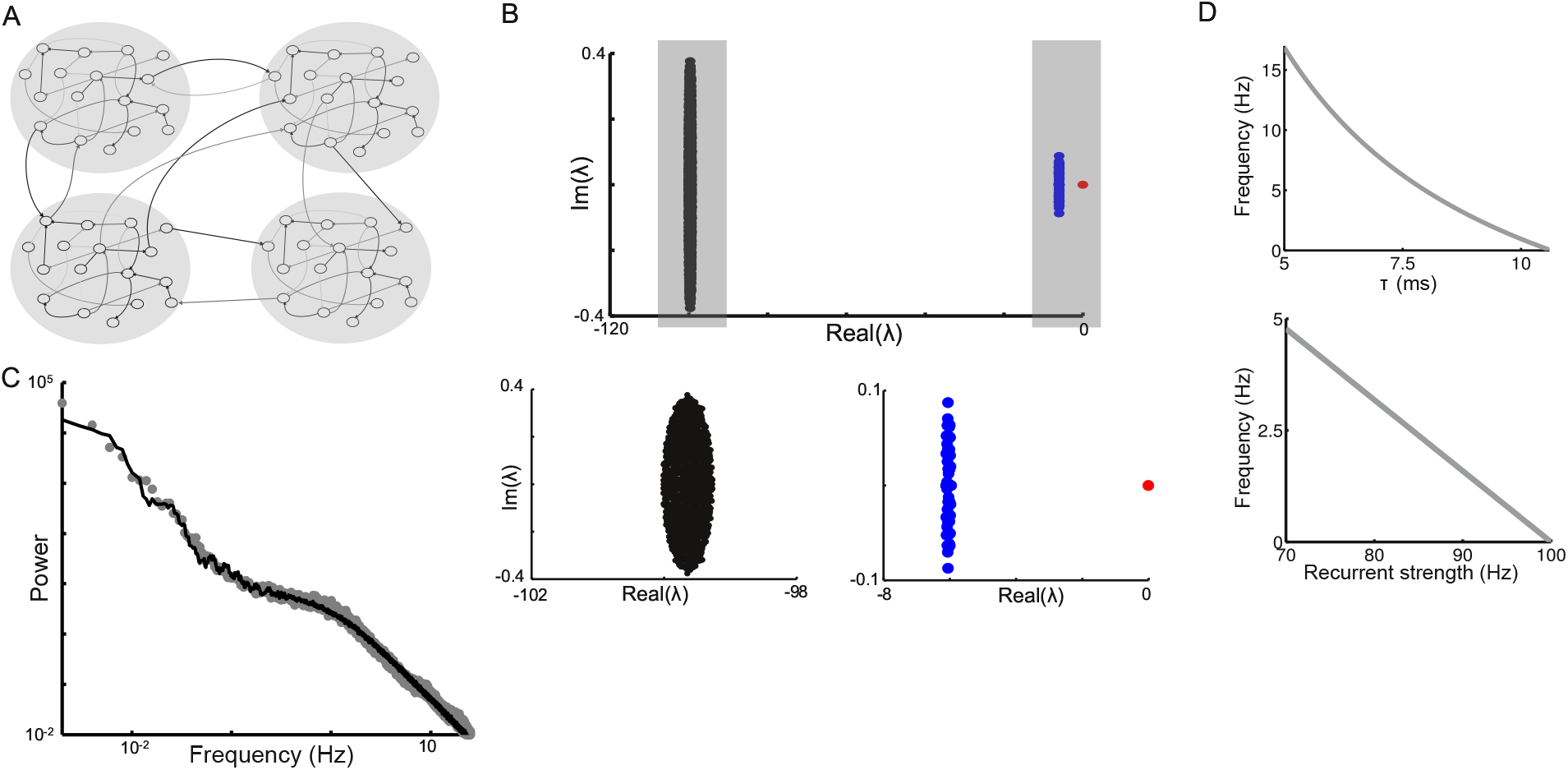
Power spectrum of a network with nodes which are themselves clusters of sub-nodes. **A**, Schematic of the network, with four clusters shown. **B**, Eigenvalue spectrum of the network. Top panel shows the full eigenvalue spectrum while the two bottom panels highlight the eigenvalues in the two gray regions. Note the hierarchical organization of the eigenvalue spectrum: the black eigenvalues reflect the timescales of individual nodes; the blue eigenvalues reflect within-cluster recurrent connections; and the red eigenvalue emerges from connections between clusters. **C**, Power spectrum of network activity (black) along with data from patient 3 (grey filled circles). **D**, Dependence of the knee frequency on intrinsic time-constant of sub-node (top panel) and on the recurrent connection strength for within-cluster connections (bottom panel). The recurrent input within a cluster is the product of connection probability, recurrent connection strength and the number of sub-nodes (i.e. size of a cluster). Increasing the intrinsic time-constant or the recurrent strength makes the network dynamics slower and thus the knee in the power spectrum shifts to lower frequencies.

In Fig. 7C we show the power spectrum of the average activity of a cluster in this network, after averaging across the individual neurons. This power spectrum is dominated by the emergent slow timescale and by the timescale of the clusters. The contribution of the very fast neural timescales is small: they originate locally and are only weakly correlated with each other and thus their contribution is strongly attenuated by averaging over the spatial scale of the cluster. Moreover, these fast timescales are on the order of milliseconds, and thus any contribution they do make is only visible at high frequencies.

As shown in Fig. 2, the knee frequencies we fit vary across electrodes and between subjects. In the model, the knee frequency corresponds to the timescale of a node or local cluster, and a variation in knee frequency suggests that the network nodes underlying each electrode show different timescales. These differences could emerge from spatial variation in the cellular and synaptic time constants of individual neurons, in the strength of recurrent interactions between the neurons and in the characteristic size of a cluster. In Fig. 7D, we show how the location of the knee frequency (i.e. the timescale of a cluster) depends on these parameters.

## Discussion

Human ECoG power spectra show evidence of simple linear dynamics dominated by a slow and a fast mode. We demonstrate how these can emerge from an underlying network with few assumptions on the connectivity: connections are random and net excitatory so as to balance the intrinsic decay of node activity. In this architecture, the fast mode reflects intrinsic properties of network nodes, and the slow mode emerges from distributed network interactions.

Fitting such recurrent networks to ECoG data reveals an eigenvalue near 0, corresponding to long correlation times. In the model, this slow timescale emerges when recurrent excitatory interactions closely balance the intrinsic decay of activity at each node (or decay plus inhibition). We find these long time-constants empirically and do not propose how recurrent excitation (or the balance between excitation and inhibition) can be driven to this point. However, several mechanisms have been proposed (Levina et al., 2007; Magnasco et al., 2009; Chialvo, 2010; Millman et al., 2010; Rubinov et al., 2011). Long correlation times are seen in systems near phase transitions (Stanley, 1999; Sethna, 2006), leading to speculation that the brain is perched at a critical point (Beggs and Plenz, 2003; Plenz and Thiagarajan, 2007; Beggs and Timme, 2012; Deco and Jirsa, 2012; Priesemann et al., 2014; Roberts et al., 2015; Bellay et al., 2015), and to suggestions that proximity to criticality provides desirable functional properties (Langton, 1990; Mitchell et al., 1993; Mora and Bialek, 2011). Our model is primarily of the system in its resting-state and we do not address these functional properties. Interestingly, both long temporal correlations and the amount of total fluctuations are suppressed upon task initiation (He et al., 2010; He, 2011, 2013; Ciuciu et al., 2012), suggesting that task performance may shift the system away from criticality (Deco and Jirsa, 2012; Fagerholm et al., 2015).

Multiple model features can be understood from the link between long timescales and recurrent excitatory interactions. The long timescales that emerge from inter-node interactions are spatially-distributed and hence correlated across nodes, while faster timescales are more local. This may explain why correlations in low-frequency activity are more widely-distributed than correlations in high-frequency activity (von Stein and Sarnthein, 2000; Buzsaki, 2006). In general, network-level activity is correlated across nodes and will become more visible after spatial averaging, such as during field potential recordings, while activity that emerges more locally will be suppressed. Thus, the model predicts that recordings that average activity over large numbers of neurons will be dominated by slower timescales (see Fig. 3E).

The clustered architecture of Fig. 7 demonstrates how the link between slow timescales and distributed interactions might operate hierarchically: neurons with fast intrinsic timescales could be arranged in clusters to produce intermediate timescales; at a higher level of organization clusters interact to produce the very long network timescales. This is compatible with recent work showing that clustered and hierarchical networks can produce dynamics with long timescales and high variability (Litwin-Kumar and Doiron, 2012; Rubinov et al., 2011). Further investigation should help identify the spatial extent of these proposed clusters.

Our model suggests potential mechanisms for the decrease in auto-correlation (as captured by the low-frequency power-law exponent ß) in ECoG and fMRI recordings upon task initiation (He et al., 2010; He, 2011; Ciuciu et al., 2012). As shown in Fig. 5, a reduction of shared inputs among nodes leads to a decrease of low-frequency power of the network fluctuations, supporting a suggestion (He, 2011) that task-induced changes may result from neurons in the local network becoming more independent (Poulet and Petersen, 2008). This decorrelation could result from nodes receiving more heterogeneous input or from an active top-down process, such as attentional decorrelation (Cohen and Maunsell, 2009; Mitchell et al., 2009).

Moreover, adding a decay of connection strength with distance caused the low-frequency power spectrum slope to become shallower. Thus, differences in connectivity structure within the same model can account for observed divergences in low-frequency power spectra. Given observations of reduced long-range effective connectivity in the human brain during slow-wave sleep (Massimini et al., 2005), this might provide a mechanistic explanation for shallower power spectra in the <0.1 Hz range during slow-wave sleep compared to the awake state (He et al., 2010). Furthermore, while a power spectrum that scales as 1/*f* is often taken to signify self-organization, we find that sharply decaying connectivity produces a spread of exponential modes; as previously shown, summing such dispersed modes can produce a spectrum that scales as 1/*f* without invoking additional physical processes (Bell, 1960; Milotti, 1995; Wagenmakers et al., 2004; Erland and Greenwood, 2007). More generally, our model suggests that the low-frequency power spectrum is sensitive to features of network organization (such as degree of averaging, connectivity decay length, excitation-inhibition balance, and correlations in the input) and could be a probe of network reconfiguration.

The average empirical power spectra (Fig. 1C) show knee frequencies at 0.49 Hz, 0.55 Hz, 0.81 Hz, 1.10 Hz and 3.47 Hz respectively, corresponding to timescales of 325 ms, 290 ms, 195 ms, 144 ms and 46 ms (timescale is 1/2*πf_fast_*). This knee frequency varies dramatically between subjects and electrodes. In the model, the frequency is determined by the timescales of individual nodes. If the nodes correspond to neurons then these might be the timescales of a slow synaptic or cellular process such as the NMDA pathway (Wang, 1999), metabotropic glutamate receptors (Zhang and Seguela, 2010), cholinergic modulation (Letzkus et al., 2011) or endocannabinoid signaling (Carter and Wang, 2007), which may vary across electrodes and subjects. For instance, time constants for synaptic transmission and single neuron dynamics may differ between sensory and association areas (Wang et al., 2008; Pereira and Wang, 2014). Alternately, the nodes might correspond to neural assemblies or clusters and the knee frequency would correspond to the timescales of these clusters, determined by local timescales and recurrent interaction within a cluster.

The location of the knee frequency shows spatial dependence and, at least in patients #1 and #3, frontal areas tend to exhibit lower frequencies (slower timescales). This may reflect a hierarchy of cortical timescales, with sensory areas processing information rapidly, whereas cognitive areas integrate information over time (Honey et al., 2012). The timescales we find are similar to those observed across cortical areas in the macaque (Murray et al., 2014), perhaps suggesting a common origin. Moreover, models suggest that a gradient of recurrent connection strengths across cortical areas could produce such a hierarchy of timescales (Chaudhuri et al., 2014, 2015). We also note that while four patients show knee frequencies near 1 Hz, patient #5 shows a faster frequency near 3.5 Hz. While our sample size is small, patient #5 is older and the difference may reflect an age-related shift in electrophysiological activity towards higher frequencies (McIntosh et al., 2010).

Above ∼ 80 Hz, ECoG power spectra have a slope steeper than 2 (Miller et al., 2009a; He et al., 2010). As observed in Miller et al. (2009a), this transition points to an even faster timescale in the data and suggests fitting the high-frequency data with a product of two Lorentzians. This produces a power spectrum that transitions from 1/*f*^2^ scaling to 1/*f*^4^ (Fig. 8). The very short timescale could emerge from fast timescales in neural input possibly imposed by synaptic time constants (Miller et al., 2009a), from a fast timescale in the output (perhaps reflecting a neuronal membrane time constant, especially in the high-conductance state (Koch et al., 1996; Destexhe et al., 2003)), or from dendritic filtering (Linden et al., 2010; Einevoll et al., 2013). Extracellular tissue filtering might also play a role (Bedard et al., 2006; Dehghani et al., 2010), although this effect remains controversial (Logothetis et al., 2007). Modeling these effects by assuming that the input to or activity from our model is convolved with a timescale on the order of milliseconds allows the model to extend into the very high frequency ranges.

**Figure 8.**
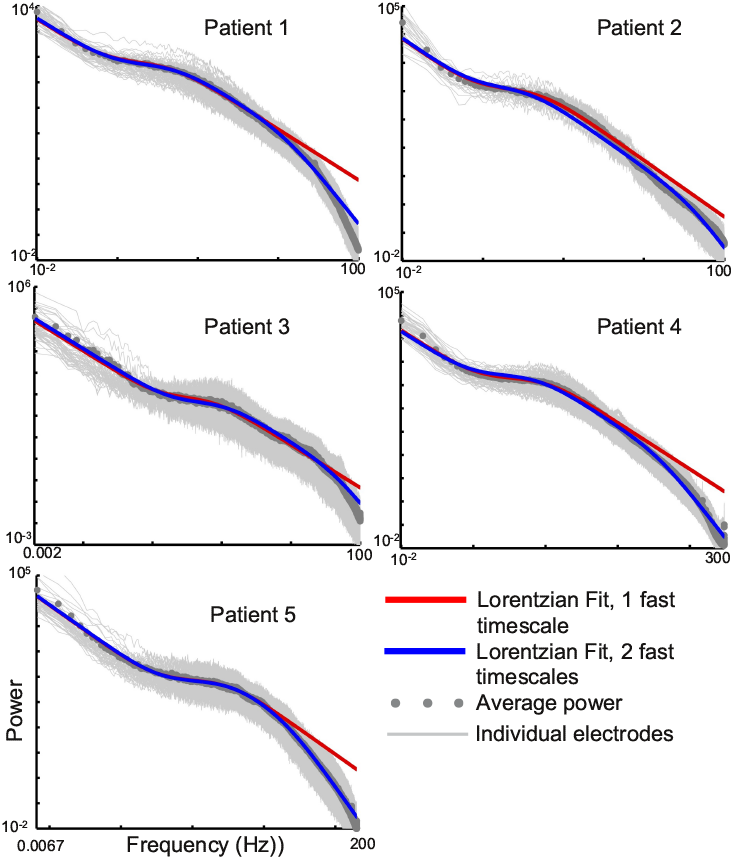
Adding a fast filtering timescale accounts for the high-frequency structure of observed power spectra. The light grey traces are recordings from each electrode while the dark grey circles are the averages across all electrodes. Red traces are the original fits with a sum of two Lorentzian functions, while blue traces are fits using a fourth free parameter, corresponding to a timescale on the order of milliseconds. The data is fit to the frequencies below 50 Hz.

While the ECoG power spectra are well-fit by linear systems, the underlying networks are likely to have non-linear components, and it will be interesting to identify neural systems with emergent macroscopic linear structure. Switches between discrete network states could produce a low frequency 1/*f*^2^ scaling, but such switches would need to be very infrequent (switching timescale close to the recording length), and the network power spectrum on shorter timescales (i.e. within a discrete state) would not show the low frequency power law scaling, in contradiction to the observations of He et al. (2010). Nevertheless rapid but infrequent shifts between states may contribute to the power spectrum, and future work should probe the differential contribution of slow drifts and abrupt switches.

In summary, our model provides a parsimonious, biologically realistic framework for interpreting broadband, arrhythmic field potentials recorded by ECoG and LFP. This framework links macroscopic arrhythmic field potentials with underlying neural network dynamics, and shows that features of the broadband power spectrum may be diagnostic of the underlying network architecture. As such, this framework may contribute to a unified understanding of previous studies and to generating and testing new hypothesis.

## Materials and Methods

*Empirical Human ECoG Data* All empirical data have been previously reported in He et al. (2010). Details of patient demographic and recording parameters can be found therein. Briefly, the study included eight patients undergoing surgical treatment for intractable epilepsy. To localize epileptogenic zones, patients underwent a craniotomy for subdural placement of electrode grids and strips followed by 1-2 weeks of continuous video and ECoG monitoring. The placement of electrodes and the duration of monitoring were determined solely by clinical considerations. All patients gave informed consent according to the procedures established by Washington University Institutional Review Board. Exclusion criteria were: (1) widespread interictal spike-and-wave discharges; (2) age < 8 years old; (3) severely impaired cognitive capability; (4) diffuse brain tissue abnormality, e.g., tuberous sclerosis, cerebral palsy; (5) limited electrode coverage.

The electrode arrays (typically 8×8, 4×5 or 2×5) and strips (typically 1×6 or 1×8) consisted of platinum electrodes of 4-mm diameter (2.3 mm exposed) with a center-to-center distance of 10 mm between adjacent electrodes (AD-TECH Medical Instrument Corporation, Racine WI).

ECoG signals were split and sent to both the clinical EEG system and a research EEG system (SynAmp2 RT, Neuroscan, DC-coupled recording). All data in the present study were from the research amplifier. Sampling rate varied from 500 to 2000 Hz across subjects, with the majority of subjects (6 out of 8) having a sampling rate of 1000 Hz.

Noisy electrodes and electrodes overlying pathologic tissue (including both the primary epileptogenic zone and areas showing active interictal discharges) were eliminated from all analyses. The remaining electrodes were re-referenced to the common average before any further analyses. Number of usable electrodes in each patient ranged from 28 to 64.

In Patients #1-#5, artifact-free, interictal-spike-free ECoG data were collected from both wakefulness and slow-wave sleep (SWS, sleep stages 3/4). Arousal state determination was based on the conjunction of ECoG and video recordings.

*The power spectrum of a linear system*. The autocorrelation function of a linear system is a weighted sum of exponentials (Gardiner, 2004). Consider a single exponential function *Ae^−λt^*, where *A* is the amplitude, *λ* is the decay constant (the time-constant is thus 1/*λ*) and *t* is time (or, more generally, the independent variable). The Fourier transform of this is 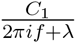, where *f* is the frequency, *i*^2^ = −1, and *C*_1_ is a normalization constant. The corresponding power spectrum is the squared magnitude: 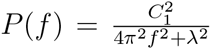 (we treat *λ* as real here). By defining *f*_0_ = *λ*/2*π* and *C*_2_ = *C*_1_/2*π*, we can rewrite this function as 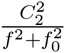. This function is a rescaled Cauchy distribution and is often called a Lorentzian.

At low frequencies (i.e. when *f* is small), the denominator is approximately 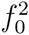 and the Lorentzian function is close to constant. On the other hand, at high frequencies (i.e. when f is much bigger than *f*_0_) this function is proportional to 1/*f*^2^. In the region where *f* ≈ *f*_0_, the function shows a transition between these two behaviors. This transition is particularly distinctive on a log-log plot, where the function shows a “knee” at *f*_0_ (see Fig. 1B), and we will refer to *f*_0_ as the characteristic or knee frequency of the Lorentzian.

*Linear fit to the data*. As depicted in Fig. 1A, the data show roughly three regimes of behavior. At very low frequencies the power spectrum is proportional to 1/*f*^2^, at intermediate frequencies the power spectrum is flat and at high frequencies the power spectrum is again proportional to 1/*f*^2^. We fit the data as the sum of two Lorentzians, with one capturing the low-frequency dependence of 1/*f*^2^ and the other capturing the intermediate flat region and the high-frequency scaling. We assume that the characteristic timescale of the low-frequency Lorentzian is longer than experimental timescales and is thus unobserved. For simplicity, we set the corresponding characteristic frequency of this Lorentzian to 0, giving us a fitting function that contains three parameters:

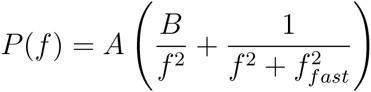

Note that, except for a small region of the power spectrum, this function is dominated by the larger of the two terms. This is depicted in the lower panel of Fig. 1B, where the sum of the two terms is well-approximated by the maximum.

In Fig. 2, we fit this functional form to the traces from each electrode. We test for spatial structure in the fitted frequencies by calculating the correlation between the frequencies of neighboring electrodes. To test for significance, we then shuffle the assignment of frequencies to electrodes and recalculate the correlation among neighbors; doing this multiple times yields a null distribution. We then compare the observed (i.e. unshuffled) correlation among neighbors with this distribution to generate a p-value.

*Power spectrum of a general linear recurrent network* We consider the general case of a linear recurrent network with *N* nodes (the nodes could be neurons or networks of neurons). The *j*th node has activity *r_j_*, which changes in time according to the equation

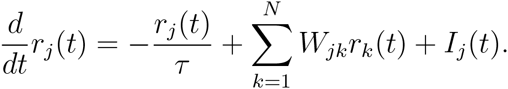

Thus, the intrinsic time-constant of a node is *τ*. The *j* th node receives input from all the other nodes (*r_k_*) with weight *W_jk_*. It also receives external input *I_j_*, which accounts for all input that does not originate within the network. In the absence of input or inter-node coupling, the firing rate of a node decays exponentially to 0 with this time-constant. Defining the firing rate vector *r(t) = [r_l_,…, r_N_]^T^* gives the equation

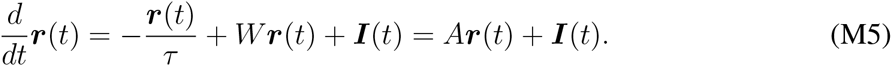

Here *W* is the matrix whose entry in row *j* and column *k* is *W_jk_*, the connection from the *k*th node to the *j*th node. *A = W* except along the diagonal, where we incorporate the intrinsic decay by subtracting 1/*τ* from each (diagonal) entry.

This system can be solved by changing into the eigenvector basis. *v_n_* is defined as a right eigenvector of *A* if *Av_n_* = *λ_n_v_n_*. Here the constant *λ_n_* is the eigenvalue corresponding to the eigenvector *v_n_*. The activity at the *j*th node is the sum of contributions from each eigenvector and (assuming that we have run the network long enough to forget initial conditions) can be written as

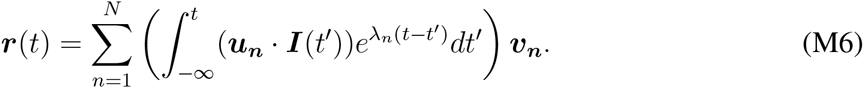

Here 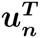 is the *n*th left eigenvector of *A*. Taking the dot product with each of the left eigenvectors allow us to convert into the eigenvector basis, so that *u_n_ · I(t’)* gives the nth component of *I(t’)* in the eigenvector basis. Thus, Eq. M6 says that to get the contribution of each eigenvector, *v_n_*, we convert the input into the eigenvector basis and filter the component along each *v_m_* with the time constant corresponding to Λ_η_. The network activity is stable if all the eigenvalues have negative real part; in this case the corresponding exponentials decay with time.

For a sub-class of matrices called normal matrices, the right and left eigenvectors are identical, and the dot product *u_n_ · I(t’)* can be replaced by *v_n_ · I(t’)*. In this case, converting into the basis given by the *v_n_* is equivalent to projecting the input along *v_n_*. Most matrices are close to normal, meaning that right and left eigenvectors tend to be aligned, and a matrix requires special structure to be highly non-normal (Trefethen and Embree, 2005). Moreover, adding randomness to the entries will tend to disrupt this special structure and make matrices more normal. Consequently, *u_n_* ≈ *v_n_*. We will use this approximation later. However, note that the connectivity matrices of networks with segregated excitatory and inhibitory populations can be quite non-normal (Murphy and Miller, 2009; Goldman, 2009).

For input with time-invariant statistics, the network dynamics are most naturally written in the Fourier basis. Taking the Fourier transform of Eq. M6 yields

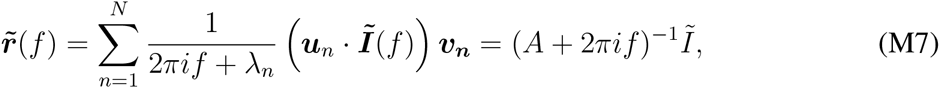

where we have defined 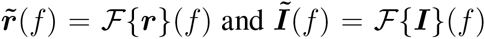 to be the vector-valued functions whose *j*th components are the Fourier transforms of the corresponding components of *r* and *I* respectively.

The power spectrum at the *j*th node is the squared magnitude of 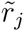. If we define

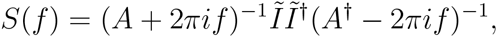

then the *j* th diagonal term corresponds to 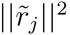. Finally, note that in the simple case when the input is uncorrelated white noise and *A* is normal, this reduces to a weighted sum of Lorentzians (this can be seen by writing 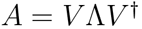, and observing that if *A* is normal then 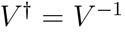).

*The random network model* We choose the connections (i.e. the entries of matrix *W*) to be sparse and random: each entry is non-zero with probability *p*, and non-zero entries are drawn from a normal distribution with positive mean *μ_conn_* and variance 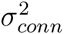 and then normalized by the size of the network (i.e. each non-zero entry 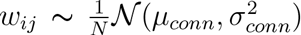). The entries thus have mean *μ = pμ_conn_/N* and variance 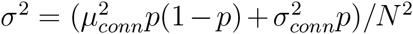 (as can be seen from an application of the law of total variance). Depending on *μ_Conn_* and *σ_conn_* this form allows a small fraction of the weights to be negative; results are similar if the normal distribution is truncated at 0. The eigenvalues of the corresponding random matrix form a cloud of points around the origin (with radius σ), along with a single outlying eigenvalue located at μ (Rajan and Abbott, 2006; Ganguli et al., 2008; Tao, 2011).

In Eq. M5, the dynamics are determined by the effective network coupling matrix *A*, whose entries are the same as that of *W* except along the diagonal, where *A_ü_ = –l/τ* + *W_ii_* (thus the diagonal entries are any self-connections minus the leak). The eigenvectors of *A* are the same as those of W, while the eigenvalues of *A* are the eigenvalues of *W* shifted by –1/*τ*. Thus the matrix *A* has a cloud of eigenvalues around *λ_fast_* = –1/*τ* and a single eigenvalue near *λ_s1ow_ = pμ_conn_*–1/*τ*.

The resulting network shows two timescales: a slow timescale with *τ_long_* = 1/*λ_slow_* = 1/(*pμ_conn_* – 1/*τ*), corresponding to the single eigenvalue near 0, and a set of fast timescales with time-constants close to τ. If the recurrent excitatory connections approximately balance the intrinsic decay then *pμ_conn_* ≈ 1/*τ*; in this case *λ_slow_* will be very close to 0, and *τ_long_* will be very long.

The eigenvector corresponding to the slow eigenvalue has special structure: it is shared across the network and approximately constant at each node (Ganguli et al., 2008). To see this heuristically, note that applying any matrix W to the constant vector [1,1,…, 1]^*T*^ yields the sum of its rows: *W*[1,1,…, 1]^*T*^ = [*Σ* _*k*_ *W*_1*k*_, *Σ* _*k*_ *W*_2*k*_,…, *Σ*_*k*_ *W*_*Nk*_]. If the rows have the same distribution, then these sums will be similar and *[Σ_k_ W*_1*k*_, *Σ_k_ W*_2*k*_,…, *Σ*_*k*_ *W*_*Nk*_] will be close to the constant vector. Thus 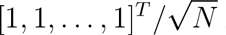 is close to an eigenvector of W (the 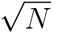 is introduced to normalize the vector).

*Power spectrum of the random network model* We now solve for the power spectrum of the network. We will use a series of approximations to analytically show that the network yields the power spectrum of Eq. 4 in the main text for uncorrelated input. In particular, we will (a) substitute the eigenvalues for the random network architecture into the expression for the Fourier spectrum; (b) use the special structure of the eigenvector corresponding to the slow mode to simplify the network response to input and (c) use the independence of the external input to each node to derive the mean power spectrum.

To begin, we substitute in the eigenvalues for this network architecture into Eq. M7, and make the approximation that all the eigenvalues in the cloud around *λ_fast_* are equal to *λ_fast_*. It then follows that

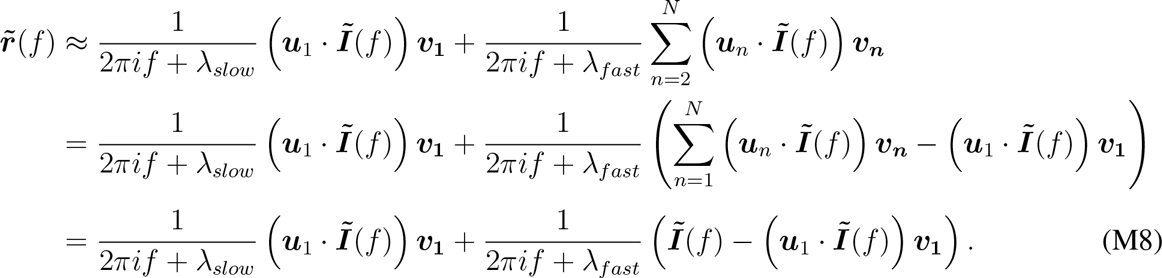

We next exploit the structure of the eigenvector corresponding to the slow mode, *v_1_* to simplify this expression. Recall that 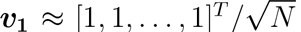 and that *u_1_* ≈ *v_1_* (this second expression is exact for a normal matrix). Substituting, we find that the power spectrum at node *j* is

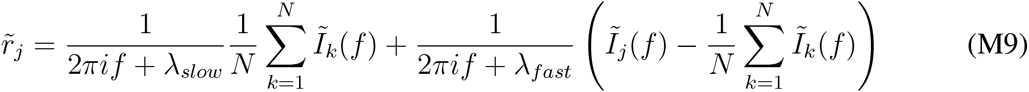

Here one factor of 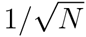 comes from *u_1_* and the other from *v_1_*. Note that 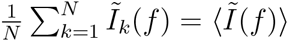, the average of 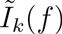 over the nodes of the network.

We now sum the activity over a fraction *α* of network nodes to produce the network activity. Note that *α* will typically be small (on the order of a few percent of network size in the fits to the data). Since the Fourier transform is linear, the Fourier transform of the summed network activity is the sum of the Fourier transformed activity at the corresponding nodes.

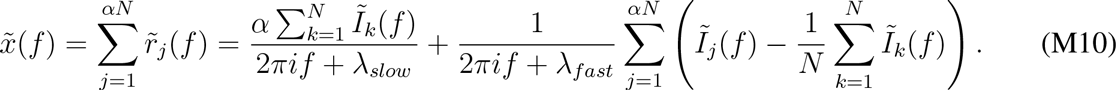

We have taken the sum over the first *αN* nodes for convenience (note that the ordering is arbitrary).

For notational convenience, wedefinethe sums 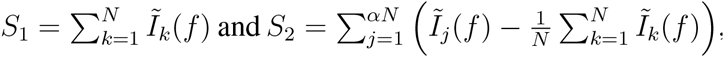, and rewrite the expression above as: 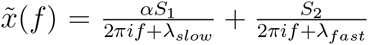. Note that the second term in *S*_2_ is independent of *j*, so the sum over *j* yields *αN* Thus 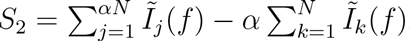.

The average power spectrum of the network activity is the expectation value of the squared magnitude of 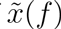.

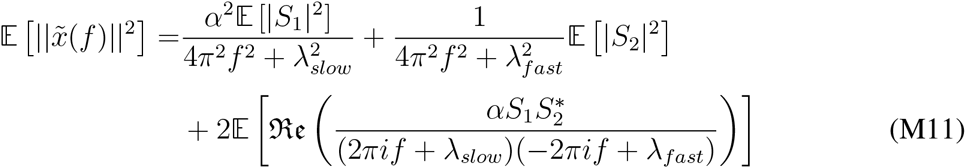

Next, note that for white noise input 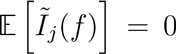 (this condition can easily be relaxed to accommodate input with a constant mean), and thus 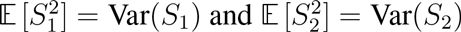.

If the input is uncorrelated across nodes and has equal variance σ^2^, then

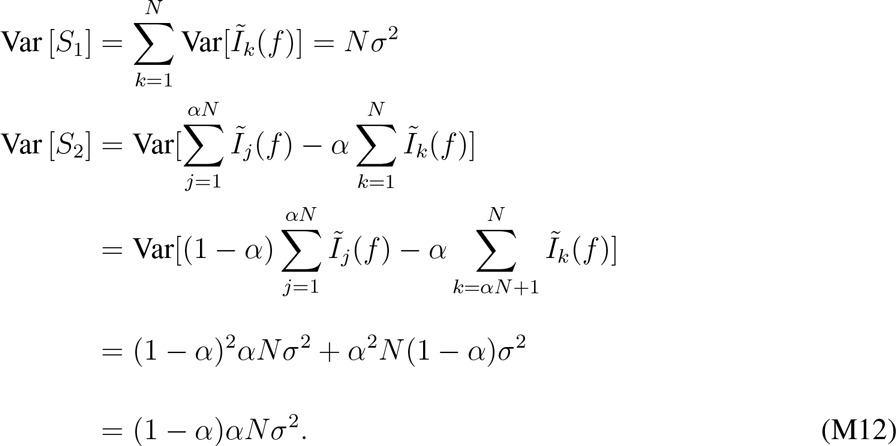

A similar analysis for the third term of Eq. M11 shows that it vanishes (at least for normal matrices where *S*_l_ and *S*_2_ are uncorrelated). Regardless, the third term is dominated by the first term at low frequencies (when *f* is smaller than any of the constants) and, when *α* is small, is dominated by the second term at high frequencies. Thus we ignore it.

Substituting the expressions for Var [*S*_1_] and Var [*S*_2_] into M11 and absorbing shared constants into an overall normalization factor, *K_norm_* then yields

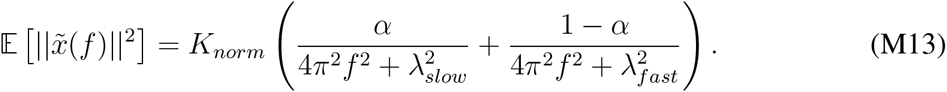

As mentioned above, this equation is valid when the input is uncorrelated across nodes. Next consider an input with some degree of correlation across nodes, consisting of a shared component with variance 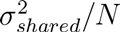 and an individual component at each node with variance 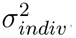. The shared component lies entirely along the slow eigenvector *v_1_* and is orthogonal to the remaining eigenvectors. Thus this expression changes to:

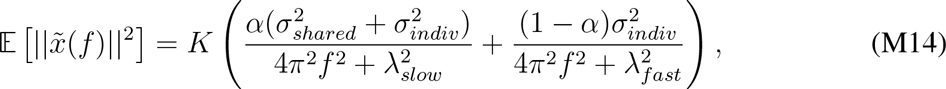

Finally, while we have considered input without temporal correlation and with comparatively simple spatial correlation, note that the analysis can be easily extended.

For Figs. 3 and 5 we set *N* = 440, *τ* = 195 ms, *μ*_*conn*_ = 25.58 Hz, *p* = 0.2 and *σ_conn_* = 2.558 Hz. In Fig. 3, we set *α* = 10/*N*, so that we average over clusters of 10 nodes. For Fig. 5 we show the average power spectrum at a single node (i.e. *α* =1 /*N*). In these simulations we set the total input variance *σ*^2^ = 1, and change the fraction of shared variance such that 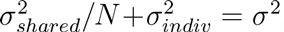. For the four simulations shown, 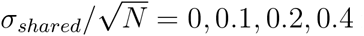 respectively.

*Network with added inhibition* For Fig. 4, we consider a network where 80% of the nodes make excitatory projections onto their targets (both excitatory and inhibitory) with mean strength *μ_Ε_* and 20% of the nodes make inhibitory projections (again, onto both excitatory and inhibitory nodes) with mean strength –*μ_I_* (where *μ_I_* > 0). This network has a similar eigenvalue spectrum to the previous network (i.e., only excitatory projections), with a cloud of eigenvalues around – 1/*τ* and a single eigenvalue corresponding to a long timescale. Now the position of this eigenvalue depends on the difference between excitation and inhibition, and is approximately located at *μ_E_p_E_* – *μ*_1_*ρ*_1_ – 1/*τ*. Here we set *N* = 440, *τ* = 195 ms, *μ_Ε_* = 51.17 Hz, *μ_ι_* = 25.59 Hz, *σ_Ε_* = 0.26 Hz, *σ_ι_* = 0.13 Hz and *p_E_* = *p_I_* = 0.2. Note that since *N_E_* = 4*N_I_*, the strength of each inhibitory connection is twice that of an excitatory connection.

*Distance-dependent connectivity* We generate the positions of the nodes randomly on a 2dimensional sheet (with hard boundary conditions, meaning that the boundary does not wrap around), with both the *x* and *y* coordinates of node positions normalized to lie between 0 and 1. Following recent experimental observations (Markov et al., 2014; Ercsey-Ravasz et al., 2013) we consider a strength of connection that decays exponentially with distance, so that *W(i, j) κ e^−A_L_d(i,j)^*, where *d(i,j)* is the distance between nodes *i* and *j*, and *λ_L_* is the inverse characteristic length of the spatial connectivity profile. Note that the results are similar if connection strength is kept fixed but the probability of a connection decays with distance, but using strength instead of probability allows us to simulate smaller networks.

The eigenvalues of this matrix can be heuristically understood as the combination of a random matrix, which produces a cloud of points around the origin, and a matrix generated from a deterministic distance-dependent connectivity profile, which produces eigenvalues scattered along the real axis between the origin and the longest timescale. The deterministic component is a translation-invariant linear operator, whose eigenvalue positions are given by the Fourier transform of the connectivity profile (Gray, 1971; Trefethen and Embree, 2005). As the connectivity profile gets more spatially-localized, the corresponding eigenvalue spectrum gets more spread out and more eigenvalues interpolate between the single slow mode and the cloud at the origin. These produce a number of intermediate frequency modes that, when smoothed together, cause a shallower slope of the power spectrum. For a sharp decay of connection strength with distance, the resulting power spectrum can show scaling of 1/*f*^−1^, as shown in Fig. 6. Note that, at least for small networks, the exact positions of the eigenvalues (and the corresponding low-frequency scaling) vary based on the distribution of distances. For the simulations of Fig. 6, we simulate networks with *N* = 100, *τ* = 195 ms and set connectivity 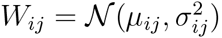. Here *μ_i,j_* = *μ*_0_*e^−λ_L_d_ij_^*, where *d* is the distance between nodes *i* and *j*, and = 0.25*μ_ij_*. For the three simulations we choose the inverse decay length *λ_L_* to be 0,10,15 and 30 respectively with corresponding *μ*_0_ = 0.77841, 1.2747, and 2.2438 Hz.

*Clustered network structure* To build the clustered network, we assume that each of the nodes in the network architecture of Figures 3 and 4 consists of *N_sub_* sub-nodes with intrinsic time-constant *τ_sub_* and within-cluster connections drawn according to 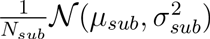. Note that this sub-network has exactly the same structure as the network shown in Fig. 3: when disconnected it has a cloud of eigenvalues around *τ_sub_* and a single eigenvalue near *μ_sub_p_sub_* – 1/*τ_sub_*. We choose this eigenvalue to give a timescale close to *τ*, where *τ* is the node time-constant from the previous section.

We then connect *N* of these sub-networks together, with sparse random connections drawn with probability *p_lr_* from a normal distribution with mean *μ_lr_*. and variance 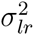. The resulting network has a large cluster of eigenvalues around *τ_sub_*, then *N* – 1 eigenvalues around *τ* and a single eigenvalue near 0, whose location is approximately *μι_r_* (*N_sub_N*)*p_lr_*.

For the simulations of Fig. 7, we set *N_sub_* = 25, *τ_sub_* = 10 ms, *μ_sub_* = 118.6 Hz, *σ_sub_* = 11.86 Hz, *N* = 44, *μ_ι_* = 25.58 Hz, *σ_ι_* = 2.558 Hz. We inject uncorrelated white-noise into all sub nodes, average the resulting activity by cluster and then plot the power spectrum of cluster activity.

*Fast timescale fits* Assuming the existence of a fast timescale in either the input to the network or the output is equivalent to convolving the network activity with an exponential with a small time-constant. In the Fourier domain, this becomes multiplication by a Lorentzian function. Thus, for Fig. 8, we fit the observed power spectra with a function of the form

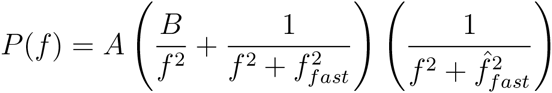

where 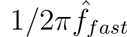 corresponds to a second fast timescale. Convolving the output of the model with a fast exponential would yield the same result

To correct for power line noise in the high-frequency region of the data, prior to fitting the power spectrum we remove data points lying within ±0.2 of 30, 60, 90, 120 and 180 Hz. Note that fits with and without filtering yield very similar parameters (not shown).

**Author contributions:** R.C., B.J.H., and X.-J.W. designed research; R.C. performed research and analyzed data; R.C., B.J.H., and X.-J.W. wrote the paper.

